# Zebrafish reveal new roles for Fam83f in hatching and the DNA damage-mediated autophagic response

**DOI:** 10.1101/2024.02.10.579757

**Authors:** Rebecca A. Jones, Fay Cooper, Gavin Kelly, David Barry, Matthew J. Renshaw, Gopal Sapkota, James C. Smith

## Abstract

The FAM83 (<u>Fam</u>ily with sequence similarity <u>83</u>) family is highly conserved in vertebrates, but little is known of the functions of these proteins beyond their association with oncogenesis. Of the family, FAM83F is of particular interest because it is the only membrane-targeted FAM83 protein. When over-expressed, FAM83F activates the canonical Wnt signalling pathway and binds to and stabilizes p53; it therefore interacts with two pathways often dysregulated in disease. Insights into gene function can often be gained by studying the roles they play during development, and here we report the generation of *fam83f* knock-out (KO) zebrafish, which we have used to study the role of Fam83f in vivo. We show that endogenous *fam83f* is most strongly expressed in the hatching gland of developing zebrafish embryos, and that *fam83f* KO embryos hatch earlier than their wild-type (WT) counterparts, despite developing at a comparable rate. We also demonstrate that *fam83f* KO embryos are more sensitive to ionizing radiation than WT embryos—an unexpected finding, bearing in mind the previously-reported ability of FAM83F to stabilize p53. Transcriptomic analysis shows that loss of *fam83f* leads to downregulation of phosphatidylinositol-3-phosphate (PI(3)P) binding proteins and impairment of cellular degradation pathways, particularly autophagy, a crucial component of the DNA damage response. Finally, we show that Fam83f protein is itself targeted to the lysosome when over-expressed in HEK293T cells, and that this localization is dependent upon a C’ terminal signal sequence. The zebrafish lines we have generated suggest that Fam83f plays an important role in autophagic/lysosomal processes, resulting in dysregulated hatching and increased sensitivity to genotoxic stress in vivo.

## Introduction

The FAM83 family is a poorly understood group of eight proteins, with members designated A-H, that is characterized by a highly-conserved DUF1669 domain (domain of unknown function 1669) at the N-termini of family members (Bartel et al., 2016, Zerbino et al., 2018, Bozatzi et al., 2018). FAM83 proteins are present in all jawed vertebrates, with no orthologs described in simpler organisms such as nematodes or *Drosophila*. There is little sequence homology outside the DUF domain, and although the DUF1669 structure is known (PDB 5LZK, Madej et al., 2014), the structures of the highly variable carboxy-termini are not.

The primary amino acid sequences of the FAM83 family give little insight into their function. The DUF1669 domain contains a pseudo-phospholipase D (PLD) domain, but no PLD enzymatic activity has been recorded for any of the family members (Cipriano et al., 2012). Until recently most attention has been paid to the oncogenic properties of the FAM83 family in numerous cancers. Broadly speaking, the FAM83 family members are found to be over-expressed in many cancers, with some (like FAM83A) being implicated in acquired therapy resistance (Lee et al., 2012). Although less is known about the role of FAM83F in oncogenesis compared with other family members, it is linked to esophageal small-cell carcinoma (Mao et al., 2016), papillary thyroid cancer (Fuziwara et al., 2019), non-small cell lung cancer (Gu et al., 2018, Fang et al., 2022, Gan et al., 2020), and breast and cervical cancer (Zhang et al., 2023, Jin et al., 2022); increased expression is also implicated in poor patient survival in other cancers (Snijders et al., 2017). Despite studies over the last decade increasingly associating the FAM83 family with oncogenesis, the mechanisms by which they drive tumorigenicity are largely unknown (Jiang et al., 2023, Yuan et al., 2022).

The FAM83 family has been associated with several signalling pathways, and members interact with many different binding partners, including c-RAF and PI3K (FAM83A, Lee et al., 2012), RAS and cRAF (FAM83B, Cipriano et al., 2013), the chromokinesin KID and dynein light chain (DYNLL1) (FAM83D, Santamaria et al., 2008, Dunsch et al., 2012), SMAD1 and CD2AP (FAM83G, Vogt et al., 2014, Cummins et al., 2017) and keratins (FAM83H, Kuga et al., 2016). Using a proteomics approach, Fulcher et al., (2018) showed that all members of the FAM83 family bind at least one isoform of casein kinase 1 (CK1), a pleiotropic and highly conserved serine/threonine kinase involved in many cellular processes and implicated in carcinogenesis (Knippschild et al., 2014). All family members bind to the CK1α isoform, and these interactions contribute to the sub-cellular localization of both proteins. This interaction with CK1α is dependent upon a F-x-x-x-F sequence motif towards the C-terminus of the DUF1669 domain of the FAM83 family proteins (Fulcher et al., 2018).

With respect to specific FAM83F interactors, both FAM83F and its most closely related family member, FAM83G, regulate canonical Wnt signaling through their interaction with CK1α, and when overexpressed in *Xenopus laevis* embryos, both induce a complete secondary axis (Bozatzi et al., 2018, Dunbar et al., 2020). Mutation of either of the phenylalanine residues in the F-x-x-x-F sequence motif of FAM83F renders the protein incapable of modulating the Wnt pathway (Dunbar et al., 2020). FAM83F also binds to and stabilizes p53 (Salama et al., 2019), and in doing so increases its activity. FAM83F was originally identified from a medaka (*Oryzias latipes*) cDNA library as one of a pool of genes that, when co-expressed with p53 in H1299 cells, leads to an increase in p53 abundance (Zhang et al., 2015). Salama et al. (2019) used similar cell culture-based methods to show that FAM83F binds to and stabilizes p53 and that the interaction between FAM83F and p53 reduced the ubiquitination of the latter. Several other putative FAM83F interactors have also been identified through mass spectrometry and have yet been explored, including Met, Ephrin B1 and 8 of the 15 human syntaxins (Gopal Sapkota, unpublished).

FAM83F is part of the ‘ignorome’ a term coined for the majority of protein coding genes that remain poorly characterized and to which there is little or no reference in the literature (Pandey et al., 2014). It is easier to study a gene that has already been studied (Stoeger et al., 2018), but to increase our understanding of biomedical science, we need to explore the roles of such proteins, particularly those implicated in human disease. These proteins are often hard to study; in the case of FAM83F, most of the experiments described in the literature are conducted in cell culture and/or by over-expression, probably because FAM83F is expressed at very low levels, and is often only detectable by immunoprecipitation (Salama et al., 2019, Dunbar et al., 2020). Insights into gene function can often be gained by studying the roles they play in development, so to elucidate the role of endogenous FAM83F in vivo, we characterized its expression in zebrafish embryos, and used CRISPR/Cas9 genome editing to knock out the primary FAM83F orthologue in zebrafish, *fam83fa*. We show that loss of *fam83fa* leads to hatching defects and impairment of cellular degradation pathways, and that *fam83fa^−/−^* embryos show increased sensitivity to genotoxic stress. We show that Fam83fa protein is itself targeted to the lysosome, making it difficult to detect at the endogenous level. We propose that Fam83fa modulates autophagic processes, including larval hatching, and that impairment of autophagy following the DNA damage response causes the increased sensitivity to genotoxic stress in *fam83fa^−/−^* embryos. To our knowledge, this is the first work investigating the role of endogenous FAM83F in the physiologically relevant in vivo context.

## Materials and Methods

### Animal housing, husbandry and microinjection

All animal work, including housing and husbandry, was undertaken in accordance with The Crick Use of Animals in Research Policy, the Animals (Scientific Procedures) Act 1986 (ASPA) implemented by the Home Office in the UK and the Animal Welfare Act 2006. Consideration was given to the ‘3Rs’ in experimental design. The zebrafish Zirc AB line was used as WT and as the genetic background for mutant line generation. Animals were maintained on a standard light/dark cycle, and for most zebrafish experiments embryos were obtained by tank mass-spawning, with males and females separated overnight and barriers removed in the morning. *Xenopus laevis* embryos were obtained by in vitro fertilization and staged according to Nieuwkoop and Faber (1994) as previously described. Embryos were maintained in Normal Amphibian Medium (Slack, 1984) until the four-cell stage, at which point they were injected with 500 pg of the indicated capped mRNA into a single ventral blastomere. Embryos were allowed to develop until approximately stage 35 at which point they were fixed in 4% paraformaldehyde. Embryos were then counted and scored for secondary axial phenotype classes as previously described (Dunbar et al., 2020).

### Generation of *fam83fa^−/−^* mutant zebrafish lines

sgRNAs were designed using CHOPCHOP (Montague et al., 2014) using the GRCz10 assembly and default search parameters. sgRNAs were selected based on their efficiency and lack of off-target effects (Table S1). The sgRNA and Cas9 protein were assembled into a ribonucleoprotein complex in vitro just prior to injection at the one-cell stage, as previously described (Burger et al., 2016, Gagnon et al., 2014, Hwang et al., 2013, Rouet et al., 2018). At 24 hours post fertilization (hpf), 10 embryos from each set of injections (sgRNA1+3 and sgRNA2+4) were screened to determine mutagenesis efficiency using standard PCR and primers designed to amplify the targeted region (Table S1). Amplicons were then TA cloned (see below) and Sanger sequenced to confirm presence of indels and to determine mutagenesis efficiency (100%). The F_0_ generation was then outcrossed to Zirc AB WT fish and a small number of embryos from the resulting clutch (F_1_) was screened for germline transmission by PCR and TA cloning as above. As germline transmission was identified in these samples, the remaining F_1_ embryos were reared to adulthood, fin-clipped and screened by PCR using the Inference of CRISPR Edits (ICE) algorithm, which allows batch analysis of CRISPR induced indels using Sanger sequencing generated .ab1 files (Hsiau et al., 2018) github.com/synthego-open/ice. F_1_ heterozygote siblings carrying identical mutations were then incrossed to generate F_2_ homozygotes, then incrossed again to generate the F_3_ maternal-zygotic (MZ-) generation, as well as being maintained as heterozygous outcrossed lines. Once the precise mutations were defined, genotyping was subsequently outsourced to Transnetyx.

### Cloning, plasmids and in vitro transcription

Total RNA was obtained from ∼50 zebrafish embryos using the NucleoSpin RNA extraction kit (Macherey-Nagel). cDNA libraries were then synthesized using the SMARTer PCR cDNA Synthesis Kit (Clontech; PT4097-1) according to the manufacturer’s instructions. ORFs for *fam83fa* and *fam83fb* were amplified from cDNA libraries and either TA cloned using the pGEM-T Easy Vector System (Promega) or the pENTR/D-TOPO vector system (Thermo Fisher). The *fam83fa^1-500^* construct was made using a reverse primer with a STOP codon at amino acid position 501 (Table S1). For microinjection, inserts were then subcloned into pCS2+ N HA tagged vectors that had been converted into Gateway (Invitrogen) destination vectors, according to the manufacturer’s instructions. PCR templates for in vitro transcription were generated from the pCS2+ Nʹ HA– tagged destination vectors. Templates were then used in an SP6 mMessage mMachine (Invitrogen) transcription reaction to generate capped mRNAs. The *hgg1* probe plasmid was a gift from Shingo Maegawa (Tanaka et al., 2018). The pDEST-CMV-N-Tandem-mCherry-EGFP plasmid was a gift from Robin Ketteler (Addgene plasmid #123216; http://n2t.net/addgene:123216; RRID:Addgene_123216) (Agrotis and Ketteler, 2015). All constructs were Sanger-sequenced by the Genomics Equipment Park Science Technology Platform at the Crick.

### Whole mount in situ hybridization (WMISH)

Digoxigenin (DIG) labelled probes were generated by in vitro transcription using a purified PCR template with the T7 promoter sequence at the 3’ end. (See Table S1 for primer sequences). A 20 µl reaction (DIG RNA labelling mix (Roche), T7 RNA Polymerase (Roche), 10X Transcription Buffer (Roche), Ribo-Lock RNase inhibitor (Thermo Fisher)) was used to generate the probe, which was then purified using the RNA Clean and Concentrator-5 kit (Zymo). Purified probes were quantified by nanodrop, checked for integrity by running on a 1% agarose gel, and stored in hybridization buffer (50% formamide, 5X SSC, 0.1% Tween 20, 50 μg/ml heparin, 500 μg/ml Torula RNA, citric acid to pH 6) at −20°C until required. WMISH was performed as previously described (Thisse and Thisse, 2008), on embryos stored in methanol at −20°C for no longer than 1 month.

### Image acquisition and live-imaging

Fixed zebrafish embryos were mounted in 2% methylcellulose on agarose coated dishes, and brightfield images captured with a Leica M165 FC microscope and a Leica DFC310 FX camera, operated by LAS V4.9 software. Live zebrafish imaging was performed using the same microscope. Embryos were imaged in glass bottom dishes in E2 medium. For confocal imaging, HEK293T cells were grown in µ-Slide 8 Well chamber slides (Ibidi). Before imaging cells were washed in PBS. Images were obtained using a Plan-Apochromat 63x/1.40 Oil DIC M27 objective of a Zeiss inverted Axio Observer LSM 710 confocal microscope, using the 405 nm, 543 nm and 639 nm laser lines to captured Z stack images. Image acquisitions and maximum intensity projections (MIP) were performed by ZEN 2010 software. Images were processed for brightness and contrast in Fiji (Schindelin et al., 2012), following the principles outlined in Rossner & Yamada (2004). For the live hatching experiment, embryos were imaged in 96-well plates, with embryos arranged into three stripes: WT embryos in columns 1-4, MZ-*fam83fa^−/−^* KO1 embryos in columns 5-8 and *MZ-fam83fa*^−/−^ KO2 embryos in columns 9-12. Over the course of three independent experiments these positions were changed to mitigate for any fluctuations in temperature at any given position. U-bottomed 96-well plates were imaged on a Ti2 inverted microscope (Nikon) with motorized XY stage (ASI) and sample temperature was maintained at 28.5°C using an environmental chamber (Okolab). Brightfield images were captured of each well every hour from 32 hpf to 80 hpf, using a 2X/0.1 NA Plan Apo objective with 1.5x intermediate magnification, with a CMOS camera (UI-3280SE, iDS) for an image pixel size of 1.15 µm. The microscope was controlled with Micro-Manager v2.0 software (Edelstein et al., 2014). For the temporal development experiment, the imaging was set up was that used in the hatching experiment and as previously described in Jones et al. (2022), with the exception of sample temperature changes where indicated.

### Ionizing radiation and MMS treatment

For ionizing radiation (IR) treatment, embryos were plated at a density of 50 per 100 mm dish in E2 medium. Embryos were treated in a GSR D1 Caesium-137 (^137^Cs) irradiator, on the shelf furthest away from the source to ensure even exposure. No more than 6 plates were treated at any one time and irradiation times were calculated using the following equation: dose required in Gy/(dose rate in Gy/min at level 3 [0.548Gy/min at 2019-2020]). Untreated embryos were taken to the irradiator alongside treated embryos but were not exposed. Following treatment, embryos were allowed to develop at 28.5°C. Embryo medium was changed at least once a day, with dead embryos and debris being removed daily. For methyl methanesulphonate (MMS) treatment, embryos were incubated in 250 µM MMS in 6-well plates, with 25 embryos per well, from shield stage (6 hpf) for 19 hours, at which point the MMS was washed off and replaced with E2 medium. Embryos were then incubated at 28.5°C and monitored until they reached 5 days post fertilization (dpf), at which point any remaining live embryos were euthanized. A minimum of 50 embryos of each genotype were used per biological replicate (n=3).

### RNA-seq

Embryos treated with ionizing radiation and control embryos were collected in 2 ml tubes, washed, resuspended in 350 μl RLT buffer (QIAGEN) and homogenized using a 1 ml syringe with a 21G x 1½” needle. Samples were snap-frozen and stored at −80°C until all three biological replicates had been collected. Samples were then subject to automated RNA extraction using the RNeasy Mini Kit (QIAGEN), with in-column DNase treatment step. RNA samples were then quantified and 25 μl of 150 ng/μl of each sample was subject to Qubit quantification and Agilent TapeStation 4200 analysis to determine quality and fragment size. After QC, ribosomal RNA (rRNA) was depleted using the Ribo-Zero Plus rRNA Depletion Kit (Ilumina), followed by library preparation using the KAPA mRNA HyperPrep Kit for Illumina Platforms, both according to the manufacturer’s instructions. Single read sequencing (75 bp) was then performed on the Illumina HiSeq 4000 platform. After filtering and trimming, Bowtie2 was used to map the sequencing reads to the GRCz11 assembly. There were a minimum of 15 million reads per condition, of which ≥6.5 million were uniquely mapped, and of these, ≥75% corresponded to exonic coding sequences. All data have been deposited in NCBI’s Gene Expression Omnibus (Edgar et al., 2002, Barrett et al., 2012) and are accessible through GEO Series accession number GSE244291 (https://www.ncbi.nlm.nih.gov/geo/query/acc.cgi?acc=GSE244291). Bioinformatic analysis was performed in R, full code is available at <u>10.5281/zenodo.10423383</u>.

### qRT-PCR

qRT-PCR primers were designed using Primer3 (http://primer3.ut.ee/) (Untergasser et al., 2012), to yield an amplicon of between 75 and 200 bp, with a T_m_ of 60°C (±2°C). Amplicons were designed to be intron-spanning, with a G or C at the 3’ end, a GC content of 40-60% to increase stability and low self-complementarity to avoid primer dimers. All qRT-PCR primers, whether designed externally or by using Primer3, were validated before use by standard PCR followed by efficiency testing by standard curve. Only primers with an efficiency of ≥90% (1.8-2.2), and a single melt curve, were taken forward for use in qRT-PCR experiments. Total RNA was extracted from pools of 20-25 embryos (unless otherwise stated) using either TRIzol chloroform extraction followed by isopropanol precipitation and LiCl precipitation or using the RNeasy Mini Kit (QIAGEN). 1 µg of RNA was then used to synthesize cDNA in a 20 µl reverse transcription reaction using 5X First Strand Buffer and MMLV-RT (Promega) with random hexamers. 10 µl qRT-PCR reactions were performed using Precision PLUS qPCR Master Mix with SYBR green (PrimerDesign) in 384-well white plates on a Roche LightCycler 480. All qRT-PCR reactions were subject to a post PCR melt-curve to validate primer specificity, and every sample, including minus-RT and no-template controls, was run in technical triplicate. The reference gene *18S* was also run on every plate in technical triplicate. Relative gene expression values were calculated using the Livak method (ΔΔCt) (Livak and Schmittgen, 2001). MIQE guidelines (Bustin et al., 2009) were followed to ensure data integrity, and all primer sequences can be found in Table S1.

### Western blotting

Zebrafish embryos were manually dechorionated and transferred to non-stick tubes in batches of 25. Liquid was aspirated off and 200 μl ice-cold 50% Ginzburg Fish Ringers solution without calcium (55mM NaCl, 1.8mM KCl, 1.25mM NaHCO_3_) was added. Embryos were pipetted up and down until the yolks came away, then a further 800 μl of Ginzburg Fish Ringers solution was added. Tubes were inverted to mix, then centrifuged at 1000g for 3 mins at 4°C. Supernatant was removed (leaving ∼20 μl) and 1 ml of ice-cold wash buffer (110mM NaCl, 3.5mM KCl, 2.7mM CaCl_2_, 10mM Tris-HCl pH 8.5) added. Tubes were shaken at 350 rpm at 4°C for 5 mins then centrifuged at 1000g for 3 mins. Cell pellets were then lysed in 100 μl/25 embryos of freshly made lysis buffer (100 μl 1M Tris pH 8.0, 20 μl 0.5M EDTA pH 8.0, 250 μl 10% IGEPAL (NP40), 100 μl 1.25M sodium β-glycerophosphate, 500 μl 1M NaF, 5 μl 20μM Calyculin A, 250 μl 200mM sodium pyrophosphate, 500 μl 10X protease inhibitor cocktail (Pierce, A32965), 3.275 ml ddH_2_O) and 4X SDS loading dye was added. Extracts were heated to 95°C for 10 mins, allowed to cool at RT for 2-3 mins, then loaded onto nUView Precast (4-20% gradient Tris-Glycine) gels (Generon) and separated by SDS-PAGE in 1X Tris-glycine running buffer (3% Tris, 14.4% glycine, 1% SDS in H_2_O). Proteins were transferred onto low fluorescence polyvinylidene fluoride (LV-PVDF) membranes in transfer buffer (3% Tris, 14.4% glycine, 0.5% SDS, 20% (v/v) methanol) using the standard “transfer sandwich” method. Following transfer, membranes were blocked in 1X Odyssey Blocking Buffer (TBS) (LI-COR) in TBS. Membranes were incubated overnight in primary antibodies in blocking buffer (anti-p53, ab77813 [9.1], (Lee et al., 2008), 1:200, anti-β-actin, (CST C4967), 1:1000, anti-HA, (Sigma H3663), 1:1000) before washing in TBST (TBS + 0.1% Tween) and overnight incubation in secondary antibodies (Goat anti-rabbit IRDye 680 RD and anti-mouse IRDye 800 CW (LI-COR), 1:10,000) in TBST with 0.02% SDS at RT. Membranes were washed, then imaged on the LI-COR Odyssey imaging system in the 700 and 800 nm channels. Images were processed for brightness and contrast in Adobe Photoshop CC (2019) following the principles outlined in Rossner & Yamada (2004). Western blotting densitometry quantification was performed by importing the .tiff file into Fiji (Schindelin et al., 2012), thresholding, analyzing particles and measuring mean gray values. Values for experimental bands were then normalized to the input bands and either plotted accordingly or expressed as a fold change value where indicated. The p53 antibody was additionally validated by western blotting protein extracts from zebrafish embryos exposed to a dose-response of IR (20, 30 and 40 Gy), to confirm increasing abundance of p53 protein (data not shown).

For *Xenopus*, embryos were obtained as described above and injected into the animal hemisphere at the one-cell stage with 500 pg of the indicated capped mRNA. Embryos were allowed to develop to stage 10 before being lysed in 10 μl/embryo ice-cold lysis buffer (1% IGEPAL, 150 mM NaCl, 10 mM HEPES, pH 7.4, 2 mM EDTA, and protease inhibitor cocktail [A32965; Pierce]). Lipids and yolk were removed from the lysate by FREON extraction (equal volume), and the aqueous phase was collected following centrifugation for 15 min at 4°C. Protein extracts were then denatured in SDS buffer before being separated by SDS–PAGE as described.

### Cell culture and tandem fluorescence assay

HEK293T cells were cultured in DMEM, high glucose (Gibco, Thermo Fisher Scientific) containing 10 % Fetal Bovine Serum (Thermo Fisher Scientific). Cells were passaged at 80 % confluency using TrypLE Express (Thermo Fisher Scientific). For transfections cells were grown in µ-Slide 8 Well chamber slides (Ibidi). Cells were transfected with a plasmid containing tandem fluorescent proteins GFP/mCherry (empty vector, EV), tandem GFP/mCherry fluorescently tagged Fam83fa protein, and tandem GFP/mCherry fluorescently tagged truncated Fam83fa^1-500^ protein (37.5 ng/well) using FuGENE HD transfection reagent as per the manufacturer’s guidelines. 24 hours following transfection, cells were treated with 50 µM Bafilomycin for 2 hours.

## Results

### *fam83fa* is expressed in the embryonic hatching gland

The zebrafish genome was interrogated to obtain the coding sequence for the open reading frame (ORF) of *fam83f*. Due to genome duplication, zebrafish frequently have more than one ortholog of a human gene (reviewed in Glasauer and Neuhauss, 2014) and *fam83f* is no exception with two orthologs, *fam83fa* and *fam83fb*, both containing 5 exons, on chromosomes 3 and 6 respectively. Fam83fa most closely aligns to FAM83F in human and other species (including mouse and *Xenopus*, Figure S1A) and contains the F-x-x-x-F sequence motif required for CK1α interaction (Fulcher et al., 2018, Dunbar et al., 2020). Furthermore, human, (and mouse and *Xenopus*) FAM83F has a C’ terminal CaaX box consisting of an invariant cysteine, two aliphatic amino acids and a terminal amino acid ‘X’. This CaaX box motif is recognised by prenylation enzymes which add either a farnesyl or geranylgeranyl moiety to the cysteine residue, targeting the protein to cell membranes (Hancock et al., 1991). Zebrafish Fam83fa, despite not having a ‘canonical’ CaaX box at the C’ terminus, is still predicted to be farnesylated in the same way as human FAM83F, on the invariant cysteine residue of the C’ terminal CIQS sequence (Xie et al., 2016, Figure S1B). By contrast, Fam83fb is 108 aa shorter than Fam83fa (447 aa compared to 555 aa) and does not contain any C’ terminal residues that are predicted to be prenylated. Moreover, whereas overexpression of Fam83fa causes a secondary axis in *Xenopus* embryos (Dunbar et al., 2020), overexpression of Fam83fb does not (Figure S1C). It is therefore likely that Fam83fa is the ortholog most closely resembling human FAM83F and we termed this the primary ortholog.

According to the White et al. (2017) RNA-seq dataset and our qRT-PCR data (Figure 1A), *fam83fa* is expressed maternally, between 2 and 4 hpf, with expression levels dropping off after zygotic genome activation (ZGA, ∼4 hpf) and peaking again around 24-48 hpf, albeit at a lower level. Whole mount in situ hybridization confirmed that *fam83fa* is expressed at late gastrula stage (90% epiboly) in the pre-polster, a distinct group of mesendodermal cells located underneath the developing forebrain from around 75% epiboly (8 hpf) (Kimmel et al., 1995, Wang et al., 2019) (Figure 1B). As this group of cells arises from the anterior axial hypoblast at around 8 hpf, this expression profile was in keeping with the qRT-PCR data, which showed that following ZGA, levels of *fam83fa* transcript start to increase after 8 hpf. The pre-polster becomes the polster by the end of gastrulation and by 12 hpf, it was visible as a U shaped mesendodermal structure in front of the boundary of the anterior neural plate (Figure 1C). The polster itself is part of the pre-chordal plate and is the rudiment to the hatching gland (Musen et al., 2012, Martínez-Romero et al., 2017, Whetzel et al., 2011). As development progresses, the polster cells migrate in an antero-dorsal direction to fan out across the yolk sac where they form the hatching gland proper (Kimmel et al., 1995, Trikić et al., 2011), and by 18 hpf *fam83fa* expression was restricted to the early hatching gland (Figure 1D). *fam83fa* expression in the hatching gland was apparent at 24 hpf and continued to persist at 48 hpf (Figure 1E-F).

**Figure 1.**
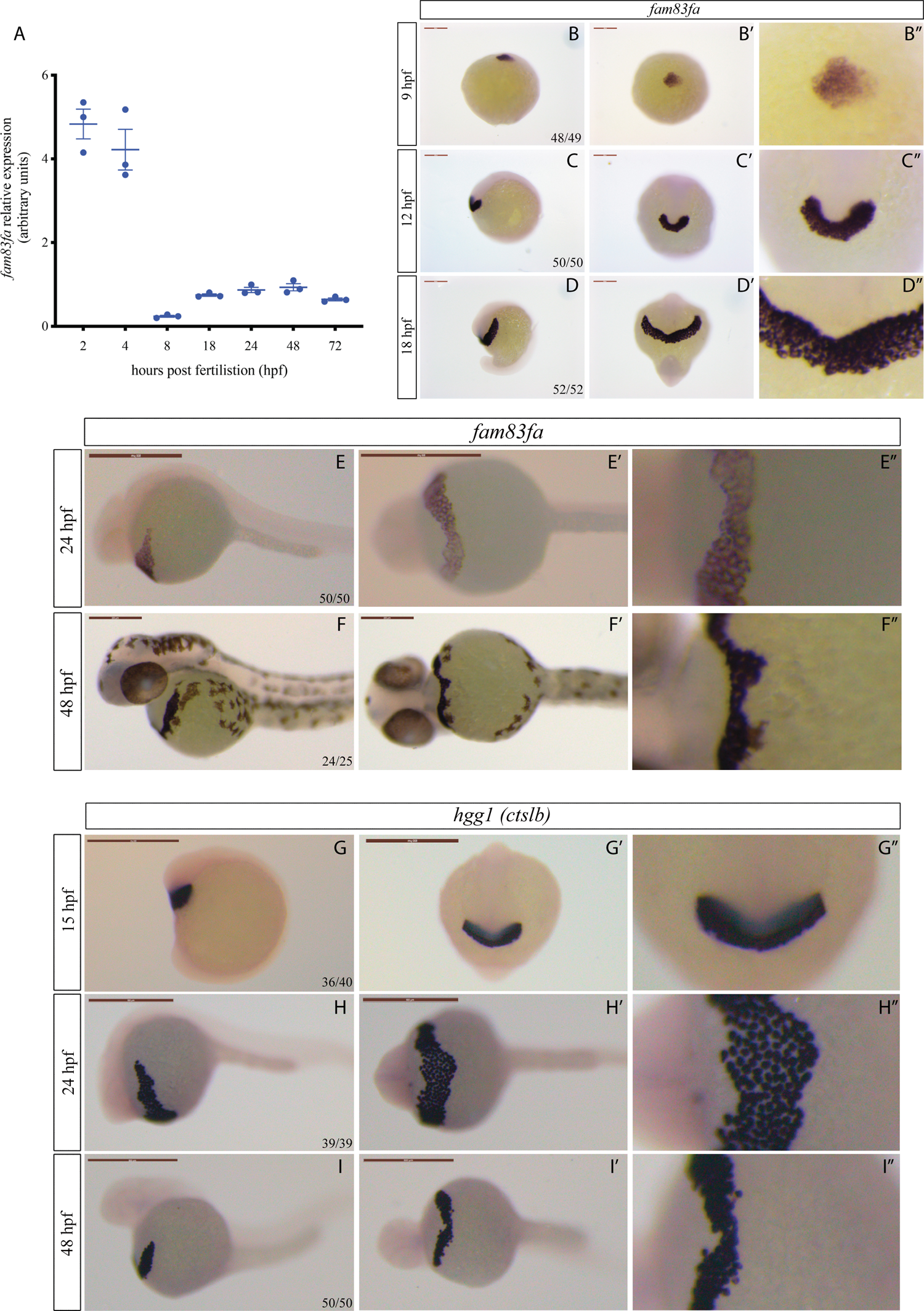
Zebrafish *fam83fa* is expressed in the hatching gland. **(A**) qRT-PCR of *fam83fa* expression (arbitrary units) in whole embryos across developmental time-series as shown. Data points = one biological replicate, n=3. Error bars = SEM. All data normalized to *18S* and calculated using the ΔΔCt (Livak) method. **(B-F)** *fam83fa* expression by whole mount in situ hybridization in stages as labeled. Lateral views (B-F), dorsal (B’) or ventral views (C’-F’) and zoomed in regions of center image (B”-F”) shown. Numbers of embryos are denoted in first image. *fam83fa* is expressed in the pre-polster, the polster and the hatching gland. Scale bars = 250 μm (except 24 hpf = 500 μm). **(G-I)** *hgg1* expression by whole mount in situ hybridization in stages as labeled. Lateral views (G-I), anterior (G’) or ventral views (H’-I’) and zoomed in regions of center image (G”-F”) shown. Scale bars = 500 μm.

The hatching gland is a specialized temporary structure present in the embryos of anuran amphibians and fish species, including teleost embryos such as zebrafish (Tatsuki et al., 2016). The primary role of the hatching gland is to release proteolytic enzymes that soften the surrounding chorion (Vogel and Gerster, 1997). This chorion softening, combined with the spontaneous movement of the larvae, allows hatching to occur, usually sometime between 48-72 hpf (Parichy et al., 2009). To validate the spatial domain of *fam83fa* expression, we performed whole mount in situ hybridization for *hatching gland gene 1* (*hgg1*), also known as *cathepsin Lb* (*ctslb*), an endopeptidase expressed in hatching gland cells at 24 hpf (Vogel and Gerster, 1997, Gardiner et al., 2005, Blanco et al., 2007). The spatial expression domain of *hgg1* matched that of *fam83fa*, confirming that *fam83fa* is expressed in the hatching gland of the zebrafish embryo (Figure 1G-I).

### Generation of *fam83fa^−/−^* zebrafish lines

Most in vivo studies into the role of FAM83F have used overexpression assays, but the best way to infer the biological function of a gene is to assess the effect of its knockout. We therefore used CRISPR/Cas9 mediated knock-out (KO) approaches to generate stable *fam83fa^−/−^* zebrafish lines.

One consideration in our KO strategy was the issue of functional redundancy, where a gene (usually a member of the same family) assumes the function of the absent gene, resulting in no apparent (or a mild) phenotype (Wagner, 1996). We also took into account the phenomenon of genetic compensation, where another gene is upregulated to compensate for the deleted gene’s function, a phenomenon that occurs in genetic mutants but not in morphants (Rossi et al., 2015). This transcriptional adaptation occurs in response to mutant mRNA decay and compensatory genes are transcriptionally upregulated in a sequence specific manner (El-Brolosy et al., 2019). With these points in mind we decided to generate four different *fam83fa^−/−^* KO lines to increase the probability that at least one line would escape these compensatory mechanisms and be more likely to show a phenotype.

Four guide RNAs (gRNAs) were designed using the online program CHOPCHOP (Montague et al., 2014) to target both transcript variants of *fam83fa* leading to complete loss of the DUF1669 domain (Figure 2A). To increase the probability that a double stranded break would be induced in the injected embryos, gRNA 1+3 RNPs and gRNAs 2+4 RNPs were injected together, to target exons 1 and 2 simultaneously. Screening of >100 animals at the F_1_ generation, using a combination of MiSeq NGS and Sanger sequencing, confirmed the high efficiency of this approach (data not shown). Of these fish, we selected four genotypes where the induced mutations were predicted to induce a premature termination codon (PTC) upstream of the F-x-x-x-F sequence motif (F275/F279), as a result of a shift in the open reading frame (ORF). These four genotypes were denoted *fam83fa^−/−^* KO1-KO4 (Table S2 and Figure 2B). As no commercial Fam83fa antibody is available, we validated our *fam83fa^−/−^* KO lines at the mRNA level. mRNAs harbouring a PTC that is 50-55 nucleotides upstream of the last exon:exon boundary trigger the non-sense mediated decay (NMD) pathway and are degraded (Hug et al., 2016). We predicted that all four of our *fam83fa^−/−^* KO lines would show reduced levels of *fam83fa* transcripts, which we confirmed by qRT-PCR (Figure 2C). All four F_3_ generation maternal-zygotic (MZ-) *fam83fa^−/−^* lines showed NMD to a similar extent, with KO1 and KO2 being most severely affected; transcripts were undetectable using a second primer set (Figure S2A). Sanger sequencing of the qRT-PCR amplicons confirmed that all predicted PTC mutations induced in the genome had been carried forward in the residual mRNA/cDNA (Figure 2D), confirming that all four MZ-*fam83fa^−/−^* KO lines were functionally null. Interestingly, qRT-PCR showed that in MZ-*fam83fa^−/−^* KO1, levels of both *fam83fa* and *fam83g* were significantly upregulated compared to both WT and the other three MZ-*fam83fa^−/−^* KO lines. This suggests that the specific mutation induced in KO1 is triggering a transcriptional adaptation response (El-Brolosy et al., 2019) that is absent in the other MZ-*fam83fa^−/−^* KO lines (Figure 2E-F).

**Figure 2.**
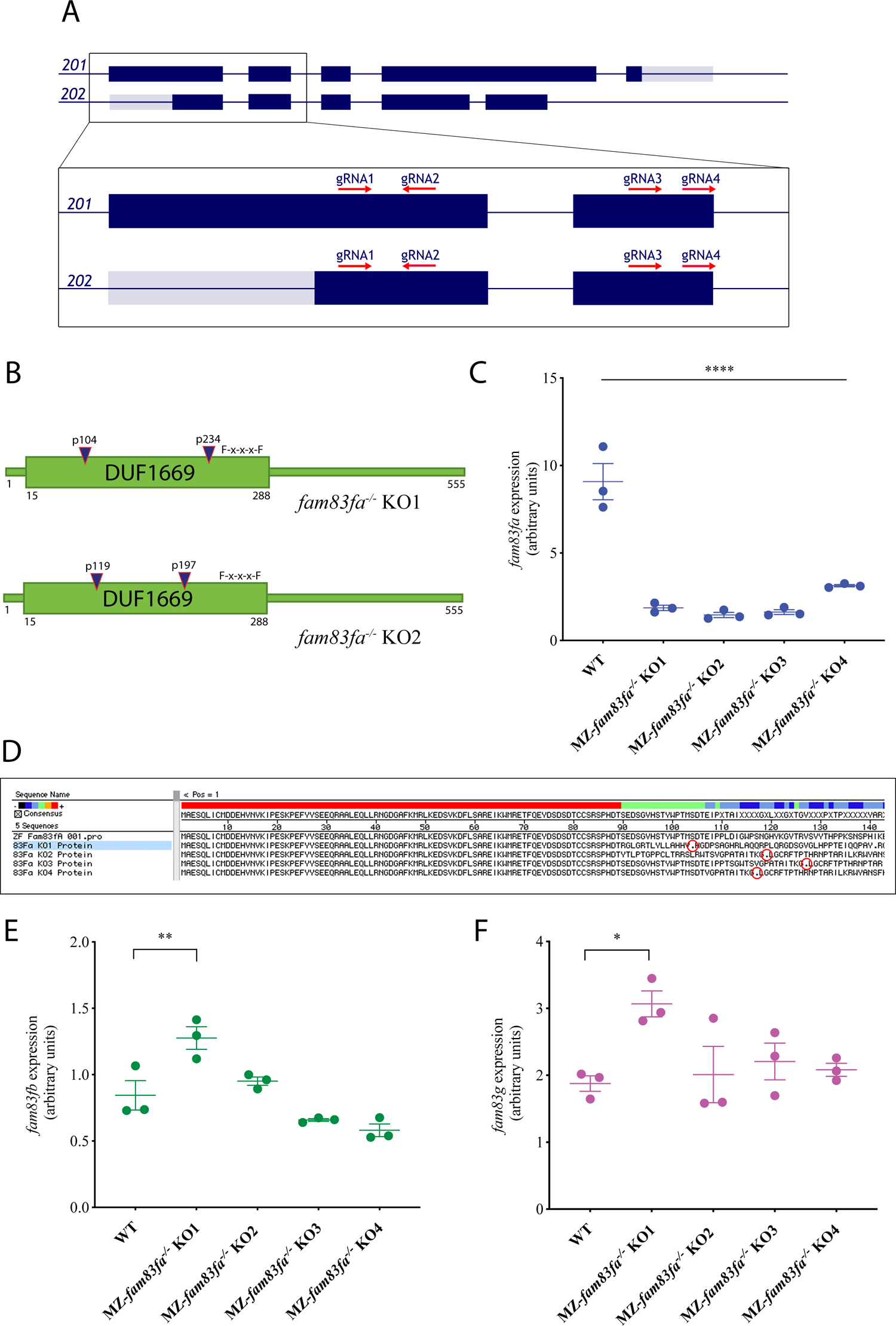
Generation of *fam83fa^−/−^* zebrafish lines by CRISPR/Cas9. **(A)** Schematic representation of *fam83fa* transcripts *201* and *202* showing CRISPR/Cas9 gRNA targeting regions. Dark blue and grey boxes denote exons and untranslated exonic regions respectively. Zoomed region shows exons 1 and 2 and the gRNA (1-4) target regions (red arrows). Transcript *201* is the presumed primary transcript; however, all gRNAs were designed to promote complete loss of the DUF1669 domain by introducing indels at the 5’ end of either transcript. **(B)** Schematic of Fam83fa showing locations of the PTCs induced in the *fam83fa^−/−^* KO1 and KO2 lines by combined injection of gRNA1 + gRNA3 (blue triangles). Exons 1 and 2 (A) code for the first 210 aa of Fam83fa, therefore all gRNAs were predicted to induce PTCs in the DUF1669 domain upstream of the F-x-x-x-F sequence motif (for simplicity, only KO1-KO2 are shown). **(C)** Relative expression of *fam83fa* in WT and MZ-*fam83fa^−/−^* KO1-KO4 homozygous embryos as labeled, normalized to *18S* (arbitrary units). Data points represent biological replicates. Error bars = SEM. ****p = <0.00005; **p = <0.005; *p = <0.05, one-way ANOVA with Tukey’s post-hoc test. **(D)** In silico translation of the Sanger sequenced qRT-PCR products amplified in (C). PTCs are denoted by red circles. Alignment performed using MegAlign software (DNASTAR). **(E)** Relative expression of *fam83fb* in WT and MZ-*fam83fa^−/−^* KO1-KO4 homozygous embryos as labeled, normalized to *18S.* Error bars = SEM. **p = <0.005; *p = <0.05, one-way ANOVA with Tukey’s post-hoc test. Data points represent biological replicates. (**F**) As (E) except showing relative expression of *fam83g*.

Having validated our *fam83fa^−/−^* KO lines, we sought to identify any phenotype associated with loss of *fam83fa.* For practical reasons, we proceeded with all subsequent experiments using two of the four MZ-*fam83fa^−/−^* lines: MZ-*fam83fa*^−/−^ KO1 and KO2. These lines were selected because of the putative presence and absence of a transcriptional adaption response, respectively.

### *MZ-fam83fa^−/−^* phenotype characterization

fam*83fa* is expressed in the hatching gland (Figure 1), so we asked whether hatching is affected in MZ-*fam83fa^−/−^* embryos. WT and MZ-*fam83fa^−/−^* KO1 and KO2 embryos were incubated at 28.5°C until they reached 24 hpf, at which point 32 embryos from each line were transferred into individual wells of a 96 well plate. Each well was maintained at 28.5°C and imaged every hour from 32 to 80 hpf. Time-lapse movies of the images captured were then manually analyzed to determine the time at which each embryo hatched. Both MZ-*fam83fa^−/−^* KO lines hatched significantly earlier than WT counterparts (Figure 3A). MZ-*fam83fa^−/−^* KO1 embryos were more severely affected, with hatching occurring on average 10 hours earlier than WT embryos, compared with 5 hours earlier for KO2 (Figure 3B). This suggests that *fam83fa* is involved in the temporal regulation of larval hatching, and that loss of *fam83fa* leads to loss of this temporal control. The primary purpose of the chorion is to protect the embryo from the external environment (Cotelli et al., 1988) and therefore the length of time for which the embryo remains within the chorion is crucial to its survival chances (Ninness et al., 2006).

**Figure 3.**
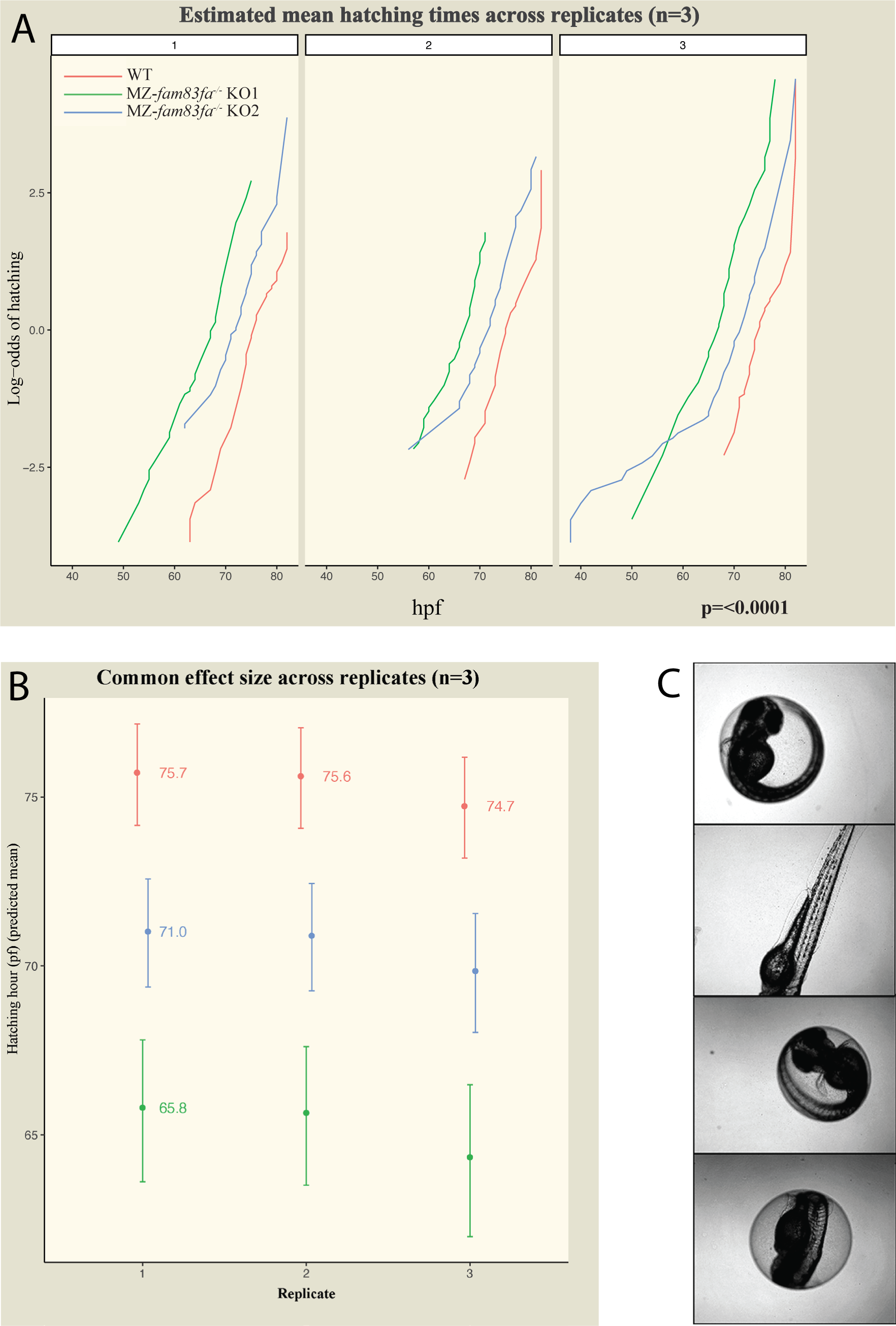
MZ-*fam83fa^−/−^* mutant embryos hatch earlier than WT. **(A)** Estimated mean hatching times across replicates (n = 3) for WT (red), MZ-*fam83fa^−/−^* KO1 (green) and MZ-*fam83fa^−/−^* KO2 (blue). Log-odds of hatching is plotted over time (hpf); where the curves cross 0.0 on the y axis, 50% of the embryos have hatched. p = <0.0001, proportional odds logistic regression. **(B)** Common effect size across replicates shown as predicted mean hatching hour. Note that MZ-*fam83fa^−/−^* KO1 embryos hatch ∼10 hours earlier than WT. **(C)** Examples of still images from time-lapse movies used to determine hatching times.

We went on to ask whether premature hatching might be secondary to a general acceleration of development. We were unable manually to detect any differences in developmental rate in MZ-*fam83fa^−/−^* embryos compared to WT, but to test this in an unbiased fashion, we used a machine-learning based object classification algorithm that enables the temporal trajectory of mutant embryos to be directly compared with their WT counterparts (Jones et al., 2022). Images were obtained of 96 embryos from WT and the most severely affected line, MZ*-fam83fa^−/−^* KO1, every 15 minutes from 4-18 hpf, then assessed by the classifier. These data confirmed there are no temporal developmental abnormalities in MZ*-fam83fa^−/−^* KO1 embryos—they develop at the same rate as WT (Figure S3). Our findings therefore demonstrate that loss of *fam83fa* in developing zebrafish embryos directly accelerates the hatching programme.

### MZ-*fam83fa^−/−^* embryos show increased sensitivity to DNA damage

With the exception of premature hatching, MZ*-fam83fa^−/−^* KO1 and KO2 embryos displayed no obvious phenotype. It is not unusual for knock-out embryos to appear normal under laboratory conditions, because some phenotypes only manifest themselves in response to specific challenges (White et al., 2013, Barbaric et al., 2007, Minchin et al., 2018, Allalou et al., 2017). With this in mind we noted that Salama et al. (2019) have demonstrated that FAM83F stabilizes p53 and promotes its activity, so we first asked whether MZ*-fam83fa^−/−^* KO embryos, in which p53 levels may be reduced, are less susceptible to cell death induced by DNA damage, as observed in *p53^−/−^* null zebrafish (Elabd et al., 2019). To this end we treated embryos from each genotype (WT, MZ*-fam83fa^−/−^* KO1 and MZ*-fam83fa^−/−^* KO2) with ionizing radiation (IR). Interestingly, MZ*-fam83fa^−/−^* KO1 and KO2 embryos were more severely affected by IR than were their WT counterparts in terms of morphological defects (bent body axis, oedema) and lethality, with MZ*-fam83fa^−/−^* KO1 again being the more severely affected (Fig S4A-B). Similar results were observed with the DNA damage-inducing agent methyl methanesulphonate (MMS) (Fig S4C-D).

The increased sensitivity of MZ*-fam83fa^−/−^* KO embryos to IR treatment suggests that Fam83f does not stabilize p53 in the early zebrafish embryo. To corroborate this conclusion, we extracted protein from WT and IR treated MZ-*fam83fa^−/−^*mutant embryos and analyzed p53 levels by western blotting (Figure 4A-B). p53 was undetectable in untreated embryos (Figure 5A and data not shown) and, as expected, there was significant induction in all IR treated samples. However, there was no detectable difference in p53 levels in MZ-*fam83fa^−/−^*mutants compared with WT embryos at either two (t1) or ten (t2) hours following IR treatment. Similarly, qRT-PCR for *p53* and its downstream targets *mdm2*, *p21*, *puma and bax* (Elabd et al., 2019, Danilova et al., 2014) showed significant upregulation in all embryos following IR treatment, but we detected no difference between WT and MZ-*fam83fa^−/−^* mutant embryos (Figure 4C-D). These data suggest that loss of *fam83fa* does not affect the stability of p53.

**Figure 4.**
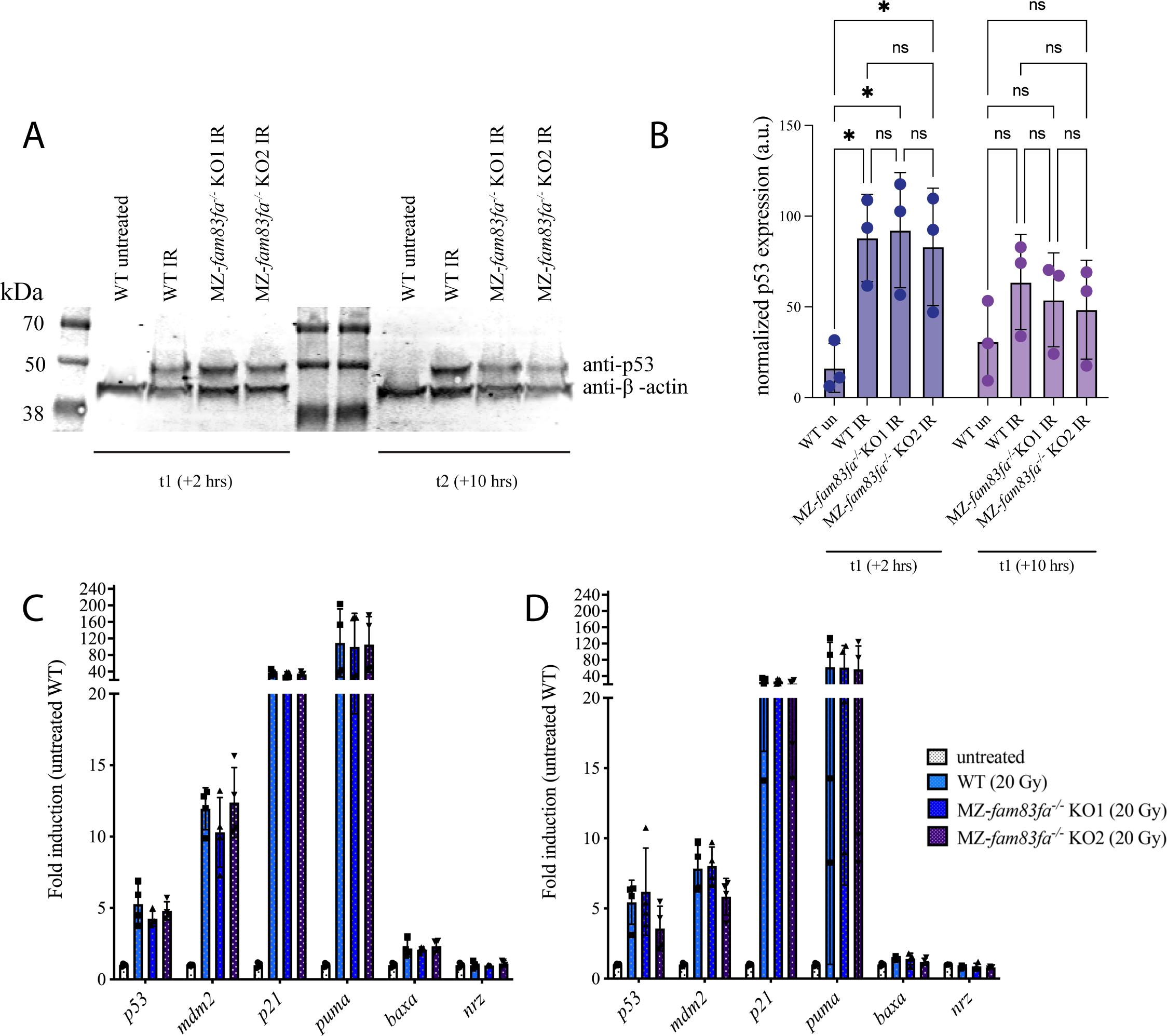
No difference in the p53 response was detected in MZ-*fam83fa^−/−^* embryos compared to WT. **(A)** Representative western blot for p53 and β-actin (input) in protein extracts from zebrafish embryos treated with IR at 24 hpf and harvested for protein extraction 2 hrs (t1) and 10 hours (t2) after treatment. **(B)** Quantification of p53 band density for 3 independent experiments as represented in (A), normalized to β-actin and expressed as arbitrary units. *p = <0.05, two-way ANOVA with Šídák’s multiple comparison test. ns = not significant. Error bars = SD. **(C)** qRT-PCR of mRNA extracted from WT or MZ-*fam83fa^−/−^* KO embryos as labeled, 2 hours following treatment with IR at 24 hpf. qRT-PCR was performed for p53 target genes, including *p53* itself, and *nrz,* the zebrafish *bcl-2* homolog. Data represent mean of four biological replicates (data-points), each consisting of 25 embryos per line, normalised to *18S* and expressed as fold induction relative to untreated WT. **(D)** As (C) except at 10 hours following IR treatment. Two-way ANOVA with Tukey’s post-hoc test showed no significant differences between IR treated WT and MZ-*fam83fa^−/−^* KO for any gene tested, Error bars = SD.

**Figure 5.**
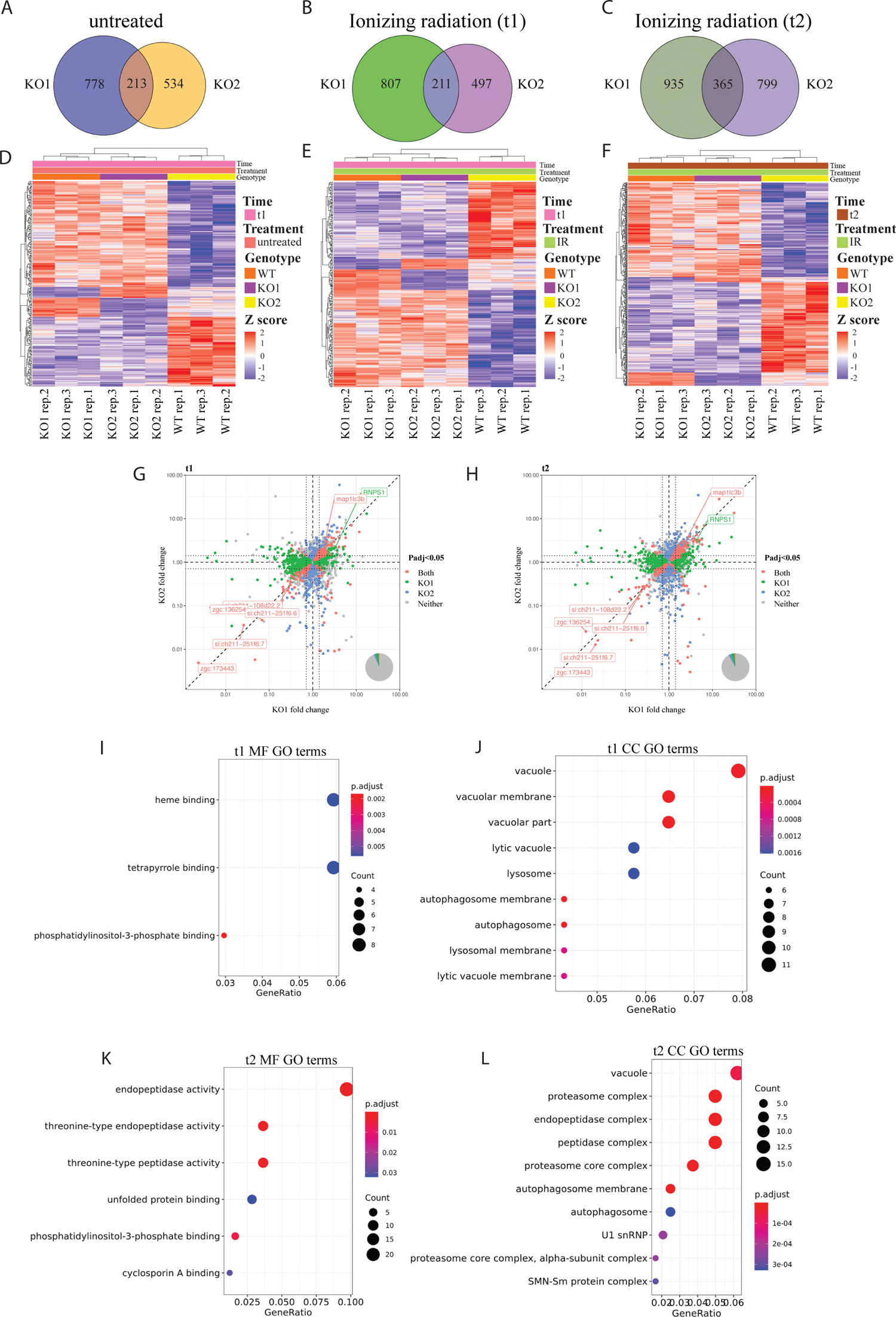
Transcriptomic analysis suggests degradation pathways are impaired in MZ-*fam83fa^−/−^* embryos. **(A)** Venn diagram showing the differentially expressed genes (DEGs) in untreated MZ-*fam83fa^−/−^* KO1 and KO2 embryos compared to WT at t1 (26 hpf). **(B)** and **(C)** As (A) except IR treated embryos at t1 (26 hpf – 2 hrs post IR treatment) and t2 (34 hpf – 10 hrs post IR treatment) respectively. **(D-F)** Heat maps of DEGs common to both KO1 and KO2 (from Venn diagram intersections above) using Euclidean distance to cluster rlog-transformed data, at conditions/times as labeled. **(G)** Volcano plot of fold change in MZ-*fam83fa^−/−^* KO1 and KO2 (*x* and *y* axes respectively) of genes with Padj <0.05, at t1. Vertical and horizontal dashed lines represent no change in KO1 or KO2 respectively, diagonal dashed line represents identical fold change in both KO1 and KO2. Dotted lines represent log2 fold change threshold of ± 0.5. Venn diagram in bottom right shows the proportion of genes in each category as shown in the legend. **(H)** As (G) except at t2. **(I)** Molecular function (MF) GO terms for DEGs common to both MZ-*fam83fa^−/−^* KO1 and KO2 at t1 (intersection of Venn diagram in (B)). **(J)** As (I) except cellular component (CC) GO terms. **(K)** Molecular function (MF) GO terms for DEGs common to both MZ-*fam83fa^−/−^* KO1 and KO2 at t2 (intersection of Venn diagram in (C)). **(L)** As (K) except cellular component (CC) GO terms.

### RNA-seq reveals downregulation of phosphatidylinositol-3-phosphate (PI(3)P) binding proteins in MZ-*fam83fa^−/−^* embryos

If not through p53 stability, why is it that MZ*-fam83fa^−/−^* embryos are more sensitive to DNA damage? To address this question, we performed bulk RNA-seq on IR-treated and untreated embryos of each genotype. Total RNA was extracted from pools of 15 embryos two hours after treatment, with additional pools being harvested eight hours later, because the DNA damage response involves early and later steps (Sirbu and Cortez, 2013) (Figure S5A).

Principal component analysis showed that most variance is described by four components (Figure S5B). PC1 accounts for developmental time (26 hpf vs 34 hpf) and PC2 accounts for treatment with ionizing radiation (Figure S5C-D). PC3 appears to represent the three genotypes (WT, MZ*-fam83fa^−/−^* KO1 and KO2), which separate roughly into three horizontal lines (Figure S5E). Most intriguingly, PC4 appears to represent the effect of loss of *fam83fa* (Figure S5F). Here, MZ*-fam83fa^−/−^* KO1 and KO2 samples cluster together on one horizontal line, clearly separate from WT. The variations underpinning PC4 may therefore be direct results of the KO of *fam83fa*, and we explored this using differential gene expression analysis.

Our analysis used DESeq2 (Love et al., 2014). Only genes with an adjusted p-value (padj) less than 0.05 and a log2 fold change (log2FC) of ± 0.5 were defined as differentially expressed (DEGs). The use of two different MZ-*fam83fa^−/−^* lines allowed us to remove differentially expressed genes that are specific to just one mutant line (Figure 5A-C); DEGs that both lines have in common are more likely to be due directly to the loss of *fam83fa*, and to underpin the variance identified by PC4 (Figure S5F).

DEGs common to both MZ*-fam83fa^−/−^* KO1 and KO2 (Venn diagram intersections, Figure 5A-C) were used to make heatmaps (Figure 5D-F). Several genes were consistently upregulated or downregulated in both MZ*-fam83fa^−/−^* KO lines at both timepoints (scatter plots, Figure 5G-H). Upregulated genes include intelectins 1 and 2 (*ITLN1* and *si:ch211-194p6.10*), carbohydrate binding proteins involved in phagocytosis (Tsuji et al., 2009, Ding et al., 2019), the lipid transport gene vitellogenin 7 (*Vtg7*), and several autophagy/lysosome-related genes including *galcb*, *arl8bb* and the autophagy associated microtubule-binding protein *map1lc3b.* Downregulated genes include *si:ch211-108d22.2*, an ATP binding protein predicted to be involved in apoptosis, and *chadla,* a transmembrane protein involved in the inflammatory response. Strikingly, the most significantly downregulated genes in MZ*-fam83fa^−/−^* KO embryos across all conditions and all time points code for predicted phosphatidylinositol-3-phosphate (PI(3)P) binding proteins, including *zgc:173443* and *si:ch211-251f6.7*. PI(3)P binding proteins are also significantly downregulated in MZ*-fam83fa^−/−^* KO embryos following IR treatment, including *si:ch211-251f6.7* and *zgc:136254*. Little is known about the roles of these genes, but they are predicted to have PI(3)P binding activity and to localize to the autophagosome/lysosome membrane (Bradford et al., 2022). A recent study by Yilmaz et al. (2021) showed that several of these predicted PI(3)P binding proteins are downregulated in *vtg3* KO zebrafish eggs, in which they also observed a concomitant upregulation of *vtg7*. This observation is in line with our dataset, suggesting a functional link between these genes. Interestingly, *rnps1,* which codes for an RNA binding protein important for NMD (Mabin et al., 2018), was only upregulated in MZ*-fam83fa^−/−^* KO1, suggesting it plays a role in the transcriptional adaptation response observed in this line (Figure 5G-H and Figure 2E-F).

GO analysis of our dataset demonstrated that genes associated with lysosomes, vacuoles and autophagosomes were significantly represented two hours after IR treatment (t1), with the majority being upregulated in both MZ-*fam83fa^−/−^* KO1 and KO2 embryos compared to WT. These genes include *lamp2* (lysosomal-associated membrane protein 2), *man2b1* (mannosidase alpha 2b1) and *map1lc3b* (microtubule-associated protein 1 light chain beta 3) (Figure 5I-J), all of which are involved in lysosomal degradation pathways (Stinchi et al., 1999; reviewed in Yoshii and Mizushima, 2017) (Figure 5I-J). By 10 hrs after IR treatment (t2), GO terms showed enrichment for genes involved in endopeptidase activity and the unfolded protein response (UPR), as well as genes associated with autophagosomes, vacuoles and the proteasome (Figure 5K-L). In addition to the PI(3)P binding proteins, downregulated genes included several proteasomal subunit genes, UPR genes such as *hspa9* (heat shock protein 9), and cyclosporin A binding proteins. In contrast, numerous genes involved in endopeptidase activity were upregulated (*furina, dpp4*, *capn2a*, *casp8l2*), together with the lysosome and vacuole associated genes *lamp2*, *man2b1*, *vac14*, *galca*, *galcb*, *map1lc3b* and others. All data have been deposited in NCBI’s Gene Expression Omnibus (Edgar et a., 2002) and are accessible through GEO Series accession number GSE244291 (https://www.ncbi.nlm.nih.gov/geo/query/acc.cgi?acc=GSE244291).

The increased sensitivity of MZ-*fam83fa^−/−^* KO embryos to DNA damage might be associated with the downregulation of PI(3)P binding proteins described above. Loss of these proteins would compromise autophagy (Palamiuc et al., 2020, Li et al., 2021), which is usually activated by genotoxic stress and is cytoprotective thanks to its role in clearing unfolded or damaged proteins (Czarny et al., 2015). Impairments in autophagic pathways have been shown to render cells more vulnerable to genotoxic agents, indeed some anti-cancer therapies are based on impairment of autophagic pathways (Levy et al., 2017). Premature hatching of MZ-*fam83fa^−/−^* KO embryos might also be explained by defects in autophagy; hatching gland cells contain large granules of proteolytic enzymes, which, when secreted at hatching, break down the surrounding chorion in a one-off event (Trikić et al., 2011). Although little is known about the regulation of hatching gland enzyme release, secretion of this sort probably occurs by autophagosome/lysosome exocytosis, and several groups have shown that autophagy is linked to the secretion of lysosomal contents and secretory granules in specialized tissues (Cadwell et al., 2008, DeSelm et al., 2011). It is therefore feasible that regulation of hatching gland enzyme release may require PI(3)P and PI(3)P binding proteins, which are downregulated in MZ-*fam83fa^−/−^* KO embryos, leading to the premature hatching phenotype we observe. This hypothesis however, requires further experimental validation.

### Fam83fa is targeted to the lysosome

The phenotypic manifestations we observed caused by loss of Fam83fa, namely, accelerated hatching and an increased sensitivity to IR, could both be potentially explained by impaired autophagy as suggested by our RNA-seq data. As such, we next asked if Fam83fa is targeted to the lysosome. Previous work has shown that human and mouse FAM83F protein is targeted to the plasma membrane by a CAAX motif (Dunbar et al., 2021, Fulcher et al., 2018) and although zebrafish Fam83fa lacks a traditional CAAX motif, the terminal amino acids are CIQS, where the terminal serine is predicted to be farnesylated in the same manner (GPS-Lipid 1.0 with high threshold parameters (Xie et al., 2016)). Following synthesis, many lysosomal membrane proteins, after transiting through the trans-Golgi network, are transported to the plasma membrane (reviewed in Saftig and Klumperman, 2009). These proteins then subsequently travel to the lysosome via the endocytic pathway. To determine whether Fam8fa is also targeted to the lysosome, we expressed tandem fluorescently tagged Fam83fa protein (GFP, mCherry) in HEK293T cells. When overexpressed, a tandemly tagged protein of this sort normally appears yellow, but if it is targeted to the acidic condition of the lysosome, GFP fluorescence is quenched faster than that of mCherry, so puncta appear red (Kimura et al., 2007). Using confocal imaging of live cells, we confirmed that Fam83fa protein appears as red puncta, suggesting that Fam83fa is targeted to the lysosome (Figure 6A). To confirm this conclusion, we treated cells with Bafilomycin A1 to inhibit acidification of the lysosome (Gagliardi et al., 1998). As expected, puncta were now yellow (Figure 6 A,B).

**Figure 6.**
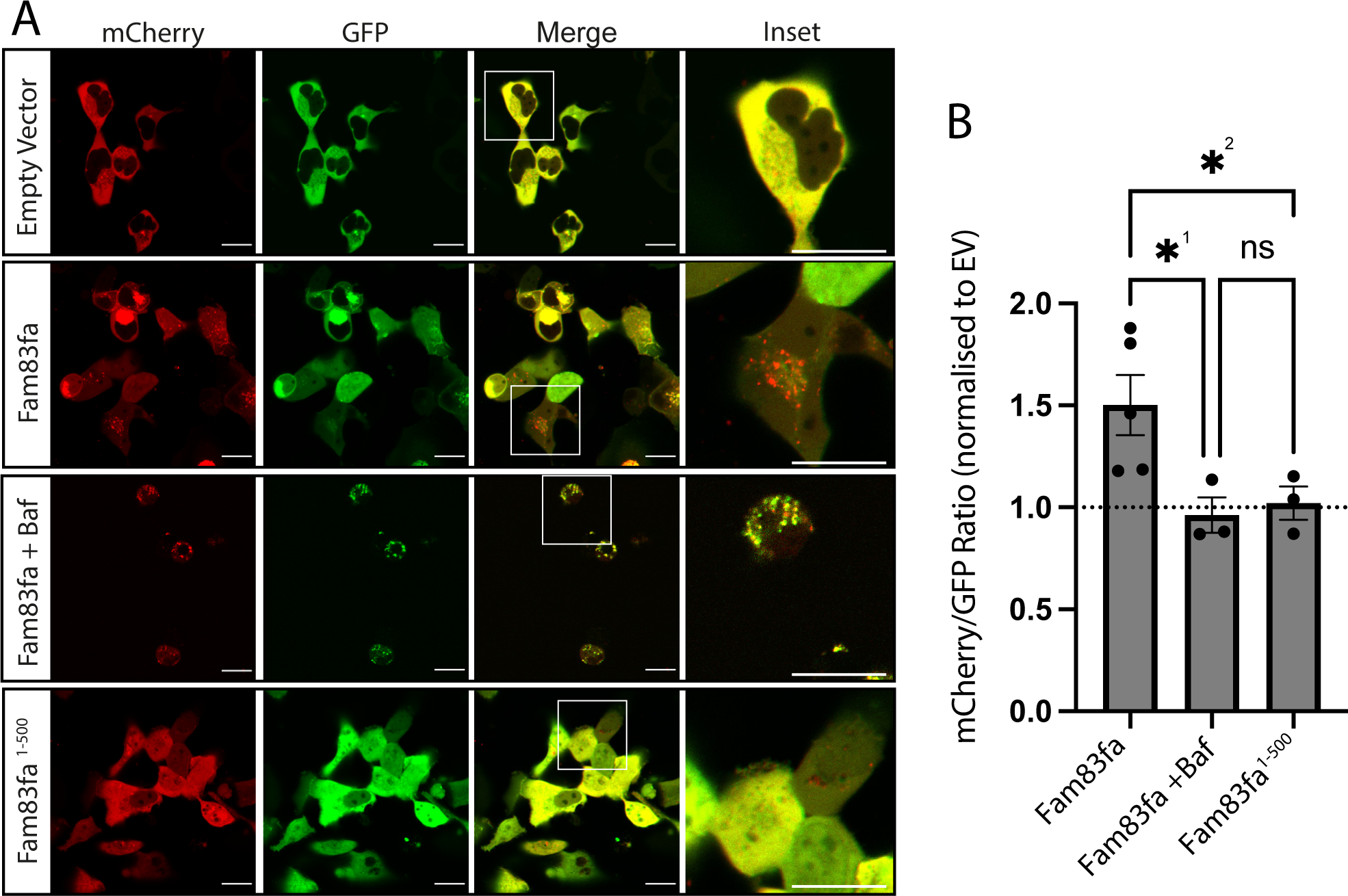
Fam83fa is targeted to the lysosome. **(A)** Representative confocal images showing HEK293T cells transfected with a plasmid containing tandem fluorescent proteins GFP/mCherry (empty vector, EV), tandem GFP/mCherry fluorescently tagged Fam83fa protein, with or without Bafilomycin A1 (Baf) and tandem GFP/mCherry fluorescently tagged truncated Fam83fa^1-500^ protein as labeled. Scale bars = 20 µm **(B)** Ratio of mCherry/GFP in all conditions shown in (A). Ratio of each biological repeat (independent transfection) is displayed normalized to empty vector control. Error bars = SEM. *^1^p = 0.0203 and * ^2^p = 0.0330, Fisher’s LSD test, ns = not significant.

Many autophagy adaptors and receptors contain LC3 interacting regions (LIR) motifs that promote their recruitment by ATG8 subfamily proteins to the lysosome, where they are subsequently degraded (Wirth et al., 2019). Using iLIR, an in silico identifier of functional LIR motifs (Kalvari et al., 2014), we identified the LIR motif DTFEFI at position 503-508 of Fam83fa. If Fam83fa protein is targeted to the lysosome for degradation upon translation as our data suggest, then this may explain why, even when overexpressed, it can be hard to detect. Western blotting experiments performed in our lab previously detected substantially more Fam83fa protein when the C’ terminal amino acids, including the predicted LIR motif, were removed (Figure S1E). To test this hypothesis, we over expressed tandem fluorescently tagged truncated Fam83fa protein (Fam83fa^1-500^) in HEK293T cells and saw complete loss of puncta, indicating that deletion of the terminal 55 amino acids of Fam83fa leads to loss of lysosomal targeting (Figure 6A,B).

## Discussion

Here we explore the function of FAM83F in the zebrafish embryo. We show that *fam83fa* is strongly expressed in the hatching gland and that MZ-*fam83fa^−/−^* KO embryos hatch earlier than their WT counterparts. We also observe that MZ-*fam83fa^−/−^* embryos are more susceptible to ionizing radiation (IR) than are their wild-type (WT) counterparts.

In attempting to understand these results, we first noted that Fam83Fa has previously been reported to stabilize p53 (Salama et al., 2019). This observation suggests that loss of Fam83fa should cause a decrease in levels of p53, and hence a reduction in IR-induced cell death—in other words, that embryos should be less susceptible to IR rather than more so, as demonstrated by *p53^−/−^* null zebrafish (Elabd et al., 2019). Our unexpected result suggested that levels of p53 might not be affected in MZ-*fam83fa^−/−^* embryos, and we confirmed this by western blotting—there were no differences in p53 levels between KO and WT embryos, whether or not they were treated with IR. Our conclusion differs from that of Salama et al. (2019), although like them we noticed that over-expression of Fam83f in the zebrafish embryo does stabilize p53 and does cause an increase in p53-mediated gene transcription and apoptotic cell death (data not shown).

With p53 unlikely to be involved directly in the Fam83F mutant phenotype, we turned to RNA-seq to compare the transcriptomes of MZ-*fam83fa^−/−^* KO and WT embryos. Our results show that the genes most significantly downregulated in MZ-*fam83fa^−/−^* mutants encode putative PI(3)P binding proteins. PI(3)P and its associated proteins play important roles in autophagy (Nishimura and Tooze, 2020, Dall’Armi et al., 2013, Palamiuc et al., 2020), the process by which unneeded cytoplasmic material is degraded and recycled via proteolytic enzymes in the lysosome. Significantly, autophagy is activated (via p53) by ionizing radiation (reviewed by White, 2016), and it can protect cells from apoptosis (Abedin et al., 2007, Muñoz-Gámez et al., 2009, Huang and Shen, 2009, Vessoni et al., 2013). It is possible, therefore, that the down-regulation of genes encoding PI(3)P binding proteins in MZ-*fam83fa^−/−^* mutants impairs autophagy and make cells more susceptible to apoptosis.

Autophagy involves the secretion of lysosomal contents and the release of secretory granules (Cadwell et al., 2008, DeSelm et al., 2011). Further evidence that Fam83F might be associated with autophagy comes from our observation that this protein is targeted to the lysosome (Figure 6A,B), which is dependent upon a signal sequence in the terminal 55 amino acids that is predicted to contain a lysosome-targeting LIR motif. Furthermore, human FAM83F interacts with syntaxins 7 and 8 (Gopal Sapkota, unpublished), which are important in late endosome:lysosome fusion (Pryor et al., 2004) and the vesicle-associated membrane protein 8 (VAMP8), which is associated with the exocytosis of secretory organelles; (Loo et al., 2009).

What of the premature hatching of MZ-*fam83fa^−/−^* embryos? Hatching gland cells contain large, lysosome-like granules of proteolytic enzymes, which, when secreted, break down the surrounding chorion in a one-off event that corresponds with hatching (Trikić et al., 2011). Little is known about how this process is timed, but the high levels of expression of *fam83fa* in the hatching gland, and the localization of the protein to the lysosome, suggest that Fam83Fa may be involved in some way, and that temporal regulation of hatching gland enzyme release is disrupted in MZ-*fam83fa^−/−^* KO embryos. This is a subject for further study, together with investigating the relationship between Fam83fa and PI(3)P binding proteins.

Using zebrafish as a model, we have uncovered a previously unappreciated role for Fam83fa in both embryonic hatching and sensitivity to DNA damage. This work provides a deeper understanding of the mechanistic role that Fam83f plays in vivo, contributing not only to our understanding of early development, but also to the pathology of human disease, given the prevalence of FAM83F dysregulation and misexpression in several human cancers.

## Acknowledgements

The authors thank Mollie Millington, Sarah Wheatley and all of the Francis Crick Institute Aquatics team for their invaluable help and patience. We thank the Francis Crick Advanced Light Microscopy Science Technology Platform (STP), the Scientific Computing STP, Dr Robert Goldstone of the Advanced Sequencing Facility (ASF) STP and Dr Nourdine Bah of the Bioinformatics and Biostatistics STP. We thank Drs Karen Vousden and Robert Ludwig (The Francis Crick Institute) for invaluable advice on the DNA damage response and Dr Sharon Tooze (The Francis Crick Institute) for autophagy advice and reagents. Finally, we thank Dr Karen Dunbar and the Sapkota lab (University of Dundee) for a most enjoyable collaboration, and members of the Smith lab for helpful discussion. This work was supported by the Francis Crick Institute which receives its core funding from Cancer Research UK (FC001-157), the UK Medical Research Council (FC001-157) and the Wellcome Trust (FC001-157).

