## Supplemental Data for "Zebrafish reveal new roles for Fam83f in hatching and the DNA damage-mediated autophagic response"

**Fig S1 Zebrafish Fam83fa is the primary FAM83F orthologue**

**(A)** Jotun Hein (PAM250) alignment of amino acids 1-300 (~DUF1669) of the different FAM83F proteins across species as shown. Zebrafish Fam83fa (fa) and Fam83fb (fb) DUF1669 domains are numbers 9 and 10 respectively. Table shows percentage identity (rows) and divergence (columns) of DUF1669 domains as labeled. Table taken from Lasergene MegAlign software (DNASTAR). **(B)** Lipid modification prediction for zebrafish Fam83fa (201 Ensembl transcript coding for the full-length protein) using GPS-Lipid 1.0 (Xie et al., 2016) with high threshold parameters. Note the high score for farnesylation, which in a 'canonical' CaaX box, is dictated by the terminal serine. **(C)** Quantification of phenotypes observed in *Xenopus* embryos injected with either 500 pg of zebrafish *fam8fa* or *fam83fb* mRNA as shown (see Dunbar et al., 2020). n = number of embryos injected, **** p=<0.0001 Chi-squared test. **(D)** Western blot of protein lysates extracted from *Xenopus* embryos injected with mRNA coding for either HA tagged zebrafish Fam83fa or Fam83fb. Blotted using anti-HA and anti-β-actin (input) antibodies. Note the robust expression of Fam83fb compared to Fam83fa. **(E)** Western blot of protein lysates extracted from *Xenopus* embryos injected with mRNA coding for either HA tagged zebrafish Fam83fa or Fam83fa$^{1\text{-}500}$. Blotted using anti-HA and anti-β-actin (input) antibodies. Note the robust expression of Fam83fa $^{1\text{-}500}$ compared to full-length Fam83fa.

**Fig S2 MZ-*fam83fa*$^{-/-}$ lines show nonsense mediated decay of *fam83fa* transcripts**

Relative expression of *fam83fa* in WT and MZ-*fam83fa*$^{-/-}$ KO1-KO4 homozygous embryos as labeled, normalized to *18S* using a second primer set (arbitrary units). Data points represent biological replicates. Error bars = SEM. ****p = <0.00005; **p = <0.005; *p = <0.05, one-way ANOVA with Tukey's post-hoc test.

**Fig S3 MZ-*fam83fa*$^{-/-}$ KO1 embryos develop at the same rate as WT**

**(A)** Scatter plot showing individual data points (images) captured every 15 minutes of each embryo (n=96 per dataset), from 4 hpf to 18 hpf. All embryos were maintained at 28.5°C except the WT 25°C plate, shown in dark blue, which acts as a control. *y* axis shows the probability that, based on appearance, the embryo is equivalent to a WT embryo at the given hpf, shown as 'predicted hpf' for simplicity. Probability calculated by object classification algorithm in ilastik, trained using WT embryos at 28.5°C (see Jones et al., 2022) **(B)** Line fit of the data in (A). Grey region around each line shows the 95% confidence interval.

**Fig S4 MZ-*fam83fa*$^{-/-}$ KO embryos are more severely affected by genotoxic agents than WT**

**(A)** Representative brightfield images of WT (left), MZ-*fam83fa*$^{-/-}$ KO1 (middle) and MZ-*fam83fa*$^{-/-}$ KO2 (right) embryos at the indicated times post IR treatment. **(B)** Kaplan-Meier curve showing survival following IR at 24 hpf of WT, MZ-*fam83fa*$^{-/-}$ KO1 and MZ-*fam83fa*$^{-/-}$ KO2 embryos (n=50) statistical analysis Log-rank (Mantel-Cox) test. **(C)** Representative brightfield images of WT (left), MZ-*fam83fa*$^{-/-}$ KO1 (middle) and MZ-*fam83fa*$^{-/-}$ KO2 (right) embryos at the indicated times post fertilization, following 19 hrs treatment with MMS as described in main text. Red arrowheads highlight embryos showing severe defects, including edema and kinked tails. **(D)** Quantification of WT, MZ-*fam83fa*$^{-/-}$ KO1 and MZ-*fam83fa*$^{-/-}$ KO2 embryos found dead at 3 or 5 dpf following MMS treatment as labeled. Each data point represents an experiment consisting of a pool of 25 embryos. ***p=<0.001, **p=<0.01 One-way ANOVA with Tukey's post-hoc. Error bars = SD. Scale bars = 1 mm.

**Fig S5 RNA-seq experimental design and principal component analysis**

**(A)** Schematic of the RNA-seq experiment characterizing the transcriptomes of WT vs MZ-*fam83fa*$^{-/-}$ KO1 and KO2 embryos following IR treatment as described in the main text. Total RNA was extracted from embryos harvested at 2 hrs (t1) and 10 hrs (t2) after treatment and sequenced on the Ilumina HiSeq 4000 platform. RNA was extracted from untreated controls at t1 only, to use for baseline comparison. Samples were collected on three separate occasions (n=3). Image created with biorender.com. **(B)** Percentage of variance in the RNA-seq dataset of the first 10 principal components. **(C)** PC2 plotted against PC1 and colored for time (t1 = 26 hpf, t2 = 34 hpf) as shown. Each dot is a sample. **(D)** As (C) except color-coded for treatment (un = untreated, IR = ionising radiation). **(E)** PC3 plotted against PC1 and colored for genotype (KO1= MZ-*fam83fa*$^{-/-}$ KO1; KO2 = MZ-*fam83fa*$^{-/-}$ KO2). **(F)** As (E) except PC4. PC4 is likely to represent variance in the data caused by *fam83fa* KO.

### Supplemental Figure S1

A

Percentage identity (upper triangle) / Divergence (lower triangle)

| | 1 | 2 | 3 | 4 | 5 | 6 | 7 | 8 | 9 | 10 | | |
|---|---|---|---|---|---|---|---|---|---|---|---|---|
| 1 | | 100.0 | 87.8 | 87.9 | 57.1 | 57.8 | 48.3 | 50.3 | 48.6 | 46.9 | 1 | Human DUF1669.pro |
| 2 | 0.0 | | 87.8 | 87.9 | 57.1 | 57.8 | 48.3 | 50.3 | 48.6 | 46.9 | 2 | Chimp DUF1669.pro |
| 3 | 13.4 | 13.4 | | 95.0 | 56.7 | 58.4 | 46.3 | 48.3 | 45.9 | 43.5 | 3 | Mouse DUF1669.pro |
| 4 | 13.3 | 13.3 | 5.2 | | 56.3 | 57.7 | 47.6 | 49.3 | 47.0 | 44.6 | 4 | Rat DUF1669.pro |
| 5 | 62.7 | 62.7 | 63.6 | 64.4 | | 90.0 | 47.0 | 47.6 | 47.3 | 44.3 | 5 | Xenopus laevis DUF1669(l).pro |
| 6 | 61.2 | 61.2 | 59.9 | 61.5 | 10.7 | | 49.0 | 48.6 | 49.3 | 44.6 | 6 | Xenopus laevis DUF1669(s).pro |
| 7 | 84.5 | 84.5 | 90.4 | 86.4 | 88.4 | 82.6 | | 84.7 | 78.5 | 62.2 | 7 | Tilapia DUF1669.pro |
| 8 | 78.9 | 78.9 | 84.5 | 81.7 | 86.4 | 83.5 | 17.1 | | 73.4 | 62.2 | 8 | Amazon Molly DUF1669.pro |
| 9 | 83.6 | 83.6 | 91.5 | 88.4 | 87.4 | 81.7 | 25.5 | 32.9 | | 63.7 | 9 | Zebrafish fa DUF1669.pro |
| 10 | 88.5 | 88.5 | 99.0 | 95.6 | 96.6 | 95.6 | 52.3 | 52.3 | 49.4 | | 10 | Zebrafish fb DUF1669.pro |
| | 1 | 2 | 3 | 4 | 5 | 6 | 7 | 8 | 9 | 10 | | |

B

| ID | Position | Peptide | Score | Cluster |
|---|---|---|---|---|
| fam83fa-201 pepti... | 552 | RRTQKKNCIQS**** | 3.861 | S-Palmitoylation: Cluster B |
| fam83fa-201 pepti... | 552 | RRTQKKNCIQS**** | 12.367 | S-Farnesylation: Non-consensus |
| fam83fa-201 pepti... | 552 | RRTQKKNCIQS**** | 3.072 | S-Geranylgeranylation: Non-consensus |

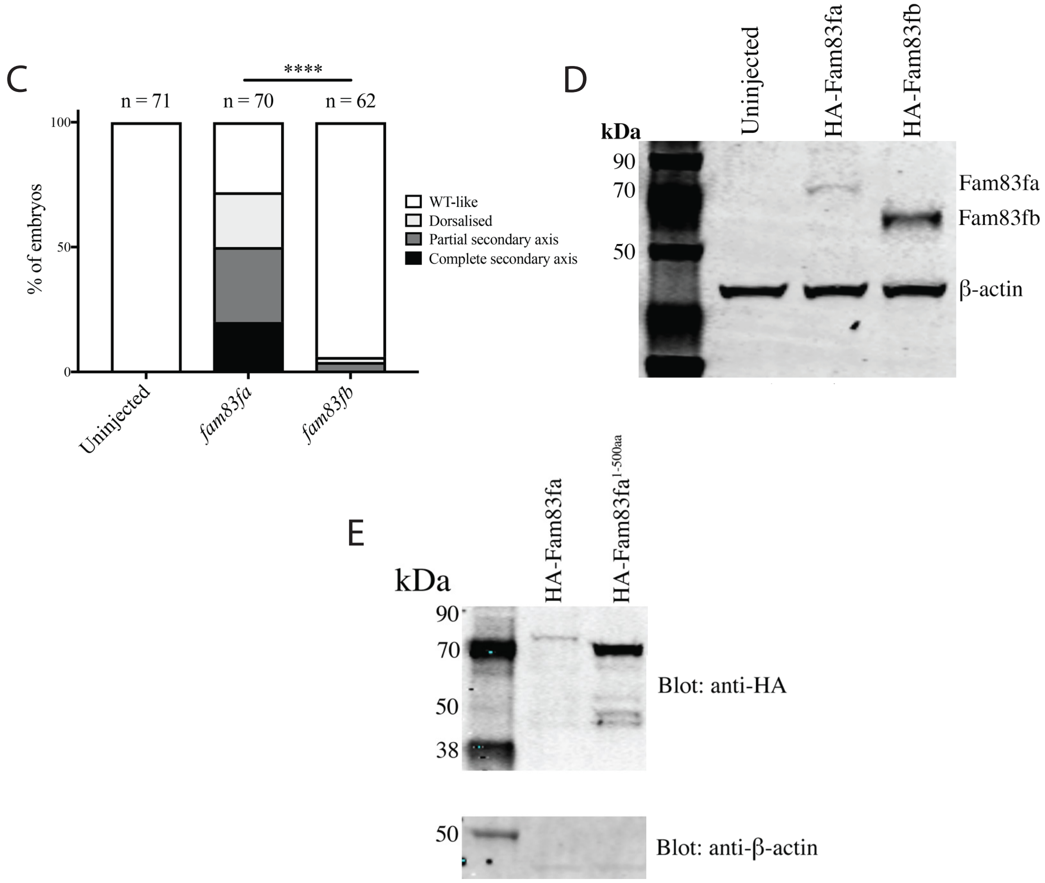

Supplemental Figure S2

A

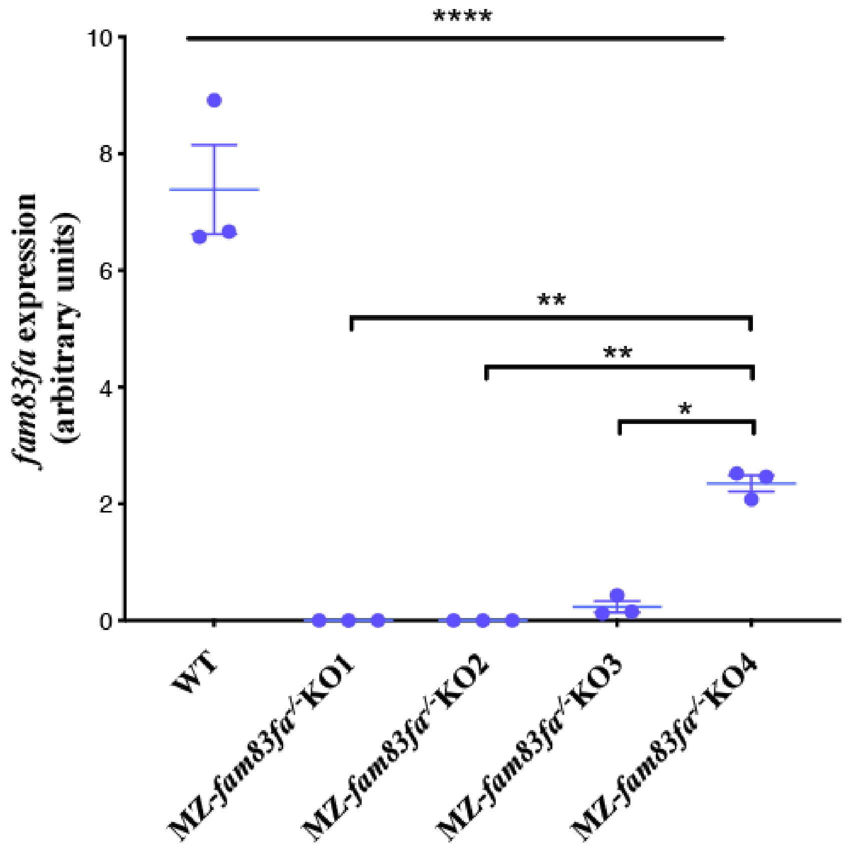

### Supplemental Figure S3

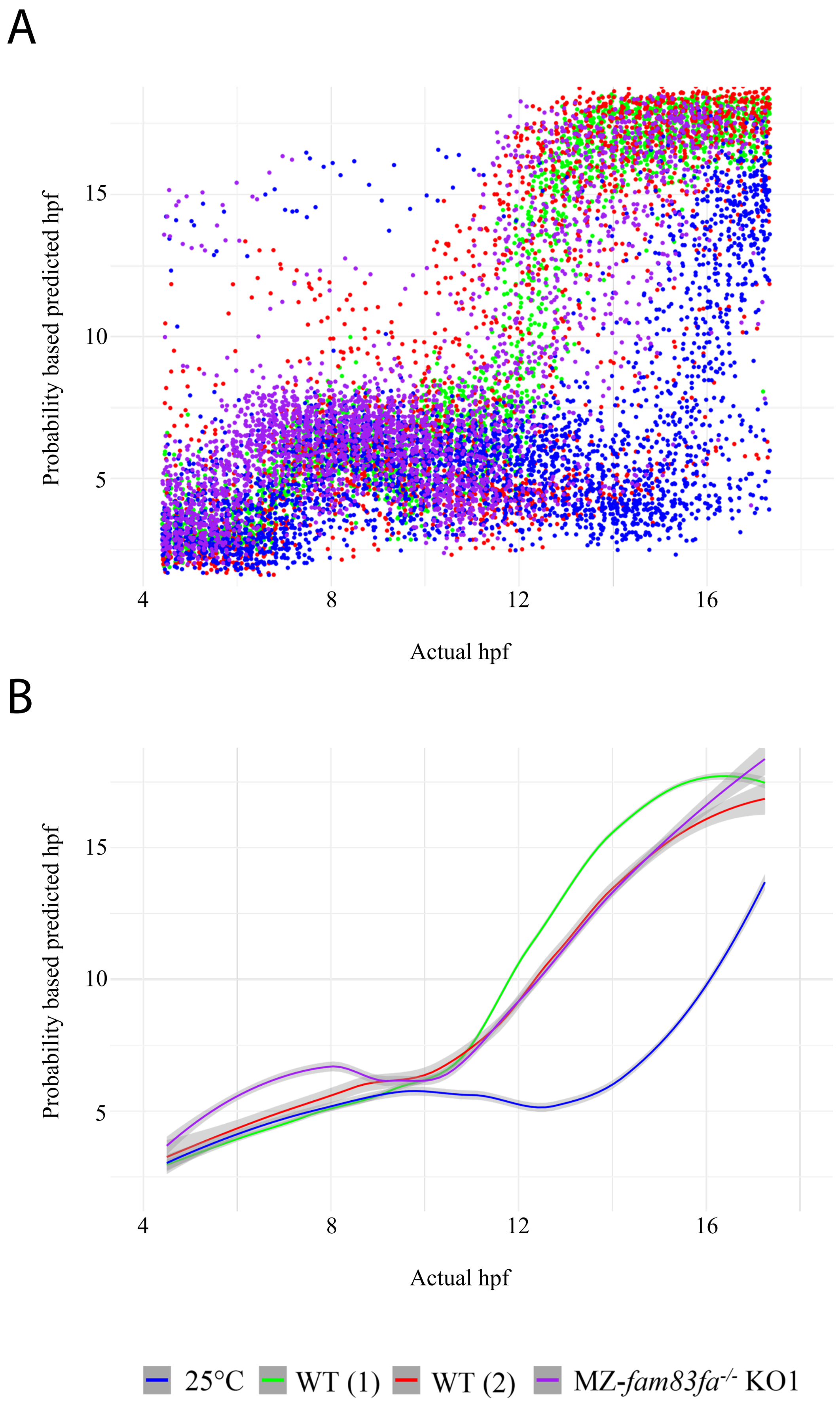

### Supplemental Figure S4

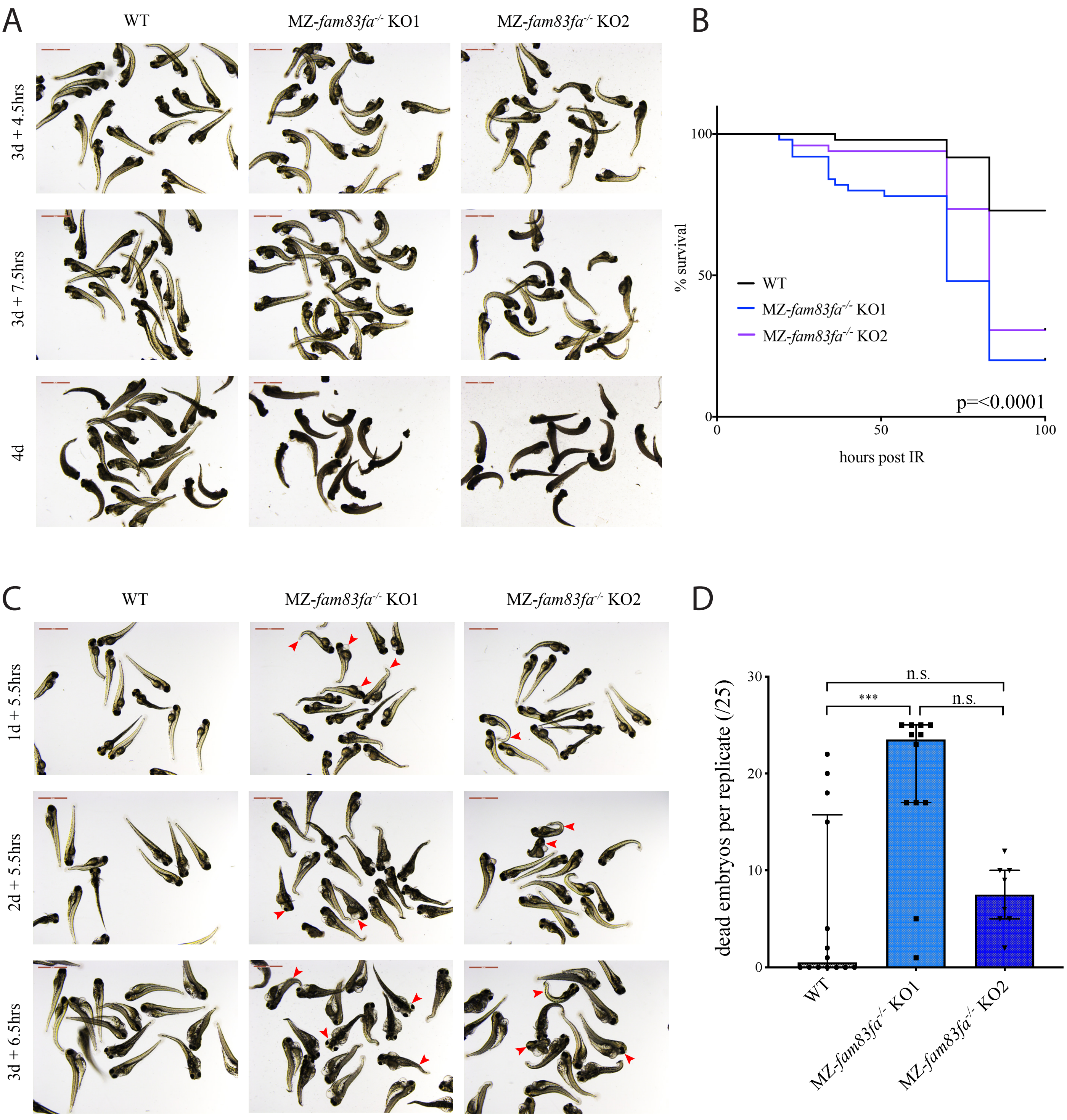

### Supplemental Figure S5

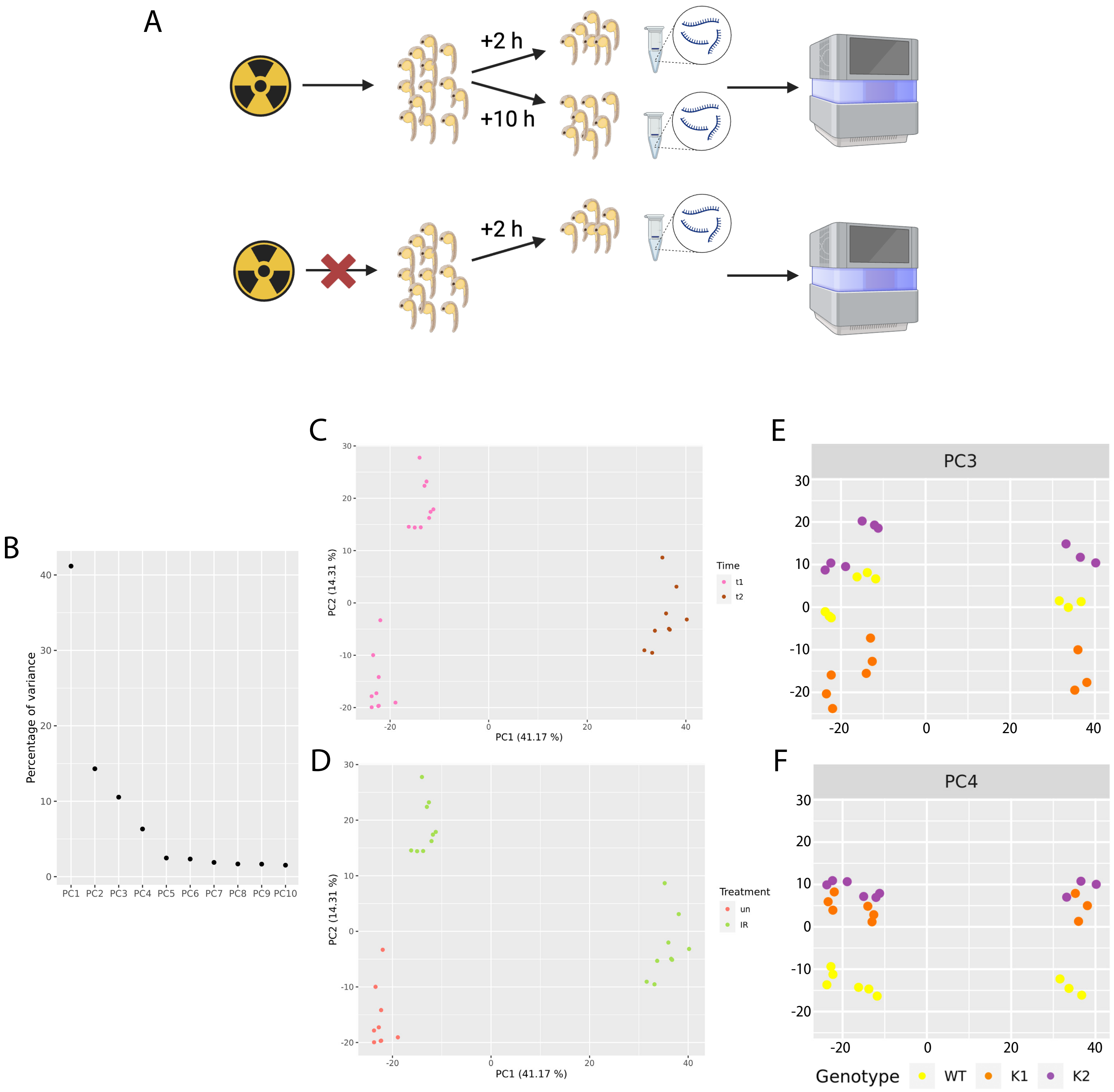
